# Maternal Time Budgets of Feral Goats at High Latitudes in Northwest Europe

**DOI:** 10.1101/2025.05.05.652180

**Authors:** R.I.M. Dunbar, Roslyn M. Anderson, Mairi E. Knight

## Abstract

Reproduction is energetically expensive for mammals, and especially so during lactation. In large-brained mammals such a primates, females invest heavily in additional feeding to fuel the costs of lactation. The evidence is more mixed in ruminant ungulates. We examine the foraging costs of lactating female feral goats living under environmentally challenging conditions in the northwest of Scotland where these effects are most likely to be exaggerated. We show, using data from three separate studies, that, compared to matched non-lactating females, lactating females do increase the time devoted to foraging, but only to a limited extent that is well below the theoretical requirements of their kids. Although they do not alter their diet, lactating females spend more time in shelter and at lower altitudes in order to reduce thermoregulatory costs. At high latitudes, the rate at which kids grow is such that females cannot afford to extend lactation much beyond two months. This will inevitably set a limit on ungulates’ capacity to produce large-brained offspring.

**Highlights:**

- Lactation is costly, obliging females to increase food intake or draw on body fat
- At high latitudes, goats increase foraging, but not enough to meet kids’ demands
- Females do not forage on richer foods, but they do try to minimise thermal costs
- As a result, females have high mortality after weaning when forage quality is poor

---

While mammalian lactation brings substantial benefit in enhancing neonatal survival rates (e.g. Power & Schulkin 2013), the costs for the mother are significant. Although the energetic costs of gestation are typically modest because the foetus is small relative to the mother (in primates, around 0.75% of the mother’s BMR, or 0.58% of her total daily energy expenditure: Ulijaszek 2002), the costs of lactation are significant. In part, this is because, after birth, the neonate increases rapidly in size (and is hence more energetically demanding) and in part because the process of lactation itself is energetically expensive. In mammals, around 20% of the additional energy (over and above the mother’s own metabolic needs) required for lactation is devoted to the physiological costs of producing milk rather than to the milk itself (Blackburn & Calloway 1976; Robbins 1983). In many herbivores, younger animals require higher protein or energy levels in their diet than adults (Arnold & Dudainski 1978), and this may impose further demands on the mother. As the infant grows, this quickly becomes a considerable burden on the mother’s foraging capacities. This inevitably has implications for how females manage their available time in order to optimise both their own and their offspring’s survival.

Increased energy demand is likely to be reflected in an increased allocation of time to feeding and greater selectivity for nutritionally high quality forage (Clutton-Brock et al. 1982a; MacWhirter 1991). Lactating red deer (*Cervus elaphus*), Stone’s sheep (*Ovis dalli stonei*), Colombian ground squirrels (*Spermophilus colombianus*), baboons (*Papio* spp.) and gelada (*Theropithecus gelada*) all spend more time feeding and foraging than non-lactating females (Altmann 1980; Clutton-Brock et al. 1982b; Seip & Brunnel 1985; Dunbar & Dunbar 1988; MacWhirter 1991). Altmann (1979) proposed that time spent feeding by lactating baboons would increase as a linear function of infant body mass, subject to the conversion efficiency of lactation (*E=*0.8: Blackburn & Calloway 1976). Her model has been tested on four different baboon populations, with broadly positive results (allowing for differences in habitat quality) (Altmann 1980; Dunbar & Dunbar 1988; Kenyatta 1995; Lycett et al. 1998).

A central assumption of the model is that an animal’s time budget is not infinitely flexible. Most species do not have significant quantities of free time that can be devoted to additional foraging when this is required by lactation (Altmann 1980; Dunbar et al. 2009). Taking time out of travel time (e.g. by moving faster between food patches) comes at the inevitable cost of increased energy expenditure, whereas taking time out of resting risks placing animals under thermal stress. In tropical habitats, for example, continuing to feed into the middle of the day when most animals go to rest means exposure to high radiant heat levels, and hence the risk of overheating (Korstjens et al. 2010); at high latitudes, the problem is more likely to be exposure to colder ambient temperatures combined with windchill, at least during winter. The time budgeting problem is exacerbated in ruminant ungulates since the bacteria responsible for gut fermentation are extremely sensitive to temperature, such that the processes of digestion automatically shut down if body temperature is elevated by any form of activity (van Soest 1994). Once ruminants have filled their gut, they have to rest in order to allow fermentation to take place so as to clear the gut before they can commence feeding again.

Ungulates differ from anthropoid primates of similar body size in that infants are weaned much earlier (6-8 weeks versus 6-8 months, respectively). Nonetheless, the increasing costs of lactation should be expressed in an increasing feeding time budget, peaking at the onset of weaning. This effect should be exacerbated in populations at high latitudes that give birth in winter because the costs of thermoregulation in low ambient temperature environments will already be placing considerable stress on females’ feeding time budgets. We explore this question in a feral goat (*Capra hircus*) population on the Isle of Rùm, NW Scotland, where the animals are exposed to very challenging climatic conditions during winter months (Dunbar & Shi 2013). Rùm is close to the northern limit at which goats can survive without being housed indoors during winter (Dunbar & Shi 2013). Most kids are born between mid-January and early February, a time of high winds and low ambient temperatures. These conditions are sufficiently taxing that females in this population hardly ever twin (despite twinning being the norm in tropical populations) and even then typically reproduce only every other year (Dunbar 2025). In addition, only 50% of newborn kids make it through the first six months of life (Gordon et al. 1987; Dunbar 2025).

We test four possible ways in which female goats might offset the costs of lactation: (1) lactating females might devote more time to foraging than non-lactating (dry) females or non-lactating (i.e. females who did not give birth that year) do, (2) the time devoted to foraging should increase linearly as a function of the energy demands of the kid as it grows (as predicted by the Altmann model), (3) lactating females should exhibit preferences for nutritionally higher quality vegetation and/or devote more time to rumination in order to exploit their ingesta more efficiently, and (4) lactating females should prefer more sheltered areas (including lower latitudes) for both foraging and resting so as to reduce thermal stress.

## Methods

### Study area and animals

The study was carried out at Harris, on the west coast of the Isle of Rùm, NW Scotland (57°0′N, 6°20′W). The goat population has been feral here since the 1770s. Female heft groups of various sizes occupied semi-exclusive territories along a 10-km stretch of coastal cliffs (Gordon et al. 1990; Dunbar 2025). Although the population was subject to intermittent culling in the 1950s and 1960s, no culling had taken place for ∼20 years prior to the start of the present series of studies in the 1990s. The Rùm goats have a main birth season between mid-January and early February; because many kids die within the first few weeks after birth, there is a secondary rut in the spring with a second birth peak in July-August (accounting for 22% of all births).

The main study site for the present studies consisted of a large, flat raised beach (Harris Bay) dominated by semi-natural *Agrostis-Festuca* grassland, with a narrow rocky foreshore; to the north and south of this the foreshore gave rise to extremely steep ∼200m sea cliffs dominated by grasses and heather (*Erica*), interspersed with rock cliffs and scree slopes. The flatter upper parts of the sea cliffs had a mixture of vegetation types (*Calluna* heath, wet heath, blanket bog, *Nardus* heath, *Schoenus* fen, *Molinia* flush, herb-rich heath and marsh). Rùm has an Atlantic climate, with high rainfall, frequent winter gales, and low but highly variable temperatures (mean monthly temperatures vary across the year between 4-14°C: Dunbar 2025). The landscape is often snow-covered in winter, especially at higher altitudes on the hilltops. Mean and minimum monthly temperatures are from Dunbar (2025); average monthly daylength is from Dunbar & Shi (2013).

The results reported here are based on studies carried out during the 1993, 2003 and 2006 winter birth seasons. Kids are suckled until they are 6-9 weeks of age (Fig. 1), after which suckling is greatly reduced. At least at this latitude, goats are only active during daylight; at night, they typically sleep in groups on the beach or in caves at beach level. During the day, the goats foraged out onto the grasslands (in Harris Bay) or up onto the cliff tops, always returning to beach level at night. Mean minimum ambient temperatures at beach level were ∼4.0°C warmer than those at an altitude of 400m asl on the adjacent hilltops, with temperatures inside the beach level caves being a further 2.0°C warmer (Dunbar & Shi 2013).

**Fig. 1.**
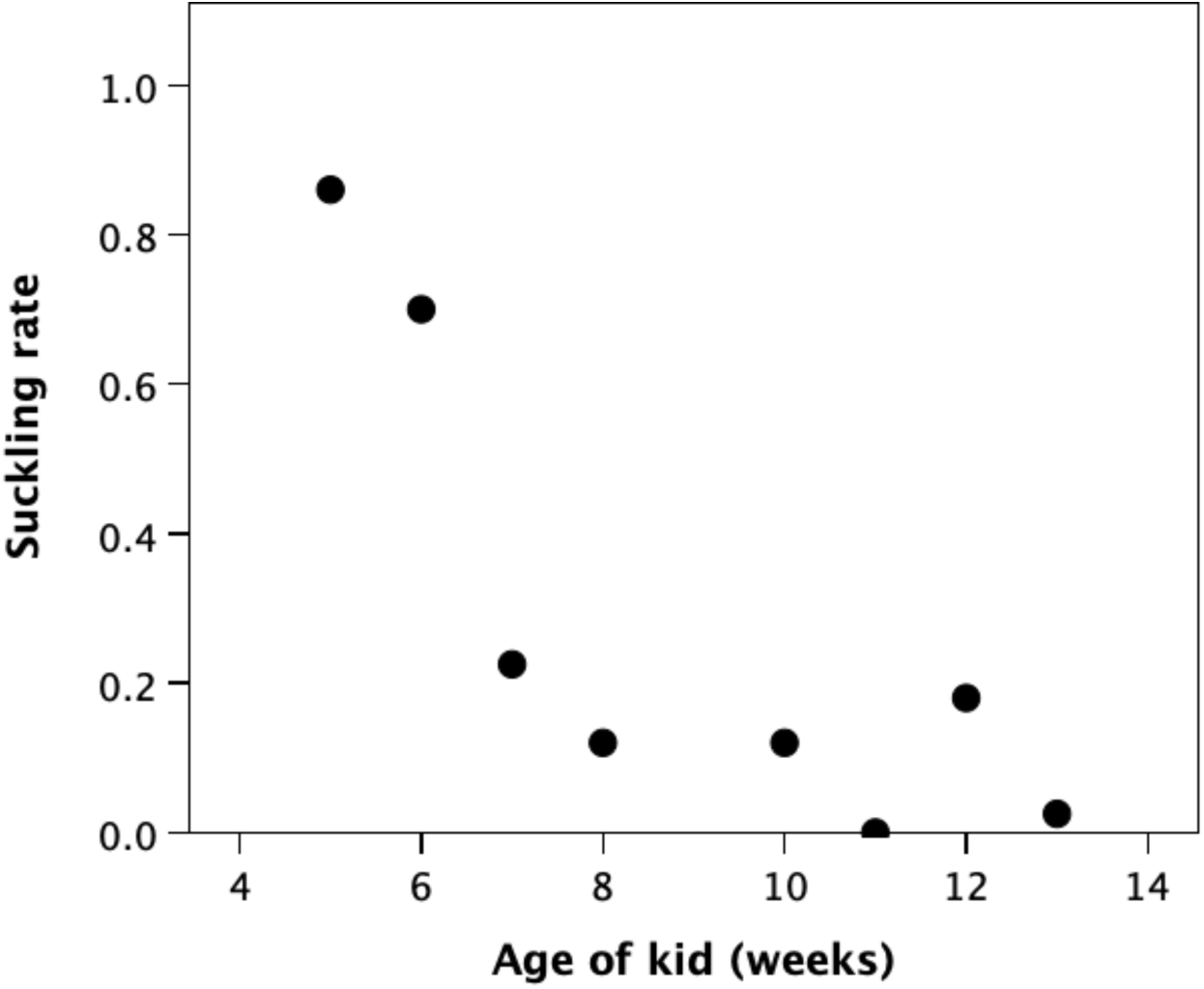
Frequency of suckling per hour by kids of different age (surviving kids only), for the 2003 birth cohort.

### Data sampling

All data were collected during focal follows of individually identified females, conducted between 08:00 and 18:00 h. Sample sizes were: six lactating and three dry females for the 1993 cohort (516 focal follows); eight lactating and four non-lactating females, plus four females whose kids died soon after birth, for the 2003 cohort (1362 focal follows); and three lactating and six non-lactating females (three of whom had given birth the previous year) in the 2006 cohort (1115 focal follows). The 1993 study was conducted on the GnP and Ruincival heft groups whose ranges lay north and south, respectively, of Harris Bay; the 2003 and 2006 studies were carried out on the Harris heft group that colonised Harris Bay in the later 1990s. Lifehistory data derive from an earlier study in 1981-2 conducted on the five heft groups occupying the sea cliffs to the north of Harris Bay (see Dunbar 2025).

Female behaviour (with kid behaviour and distance of mother to kid), location, vegetation type, whether or not in shelter, weather conditions (including wind speed on the Beaufort scale), other group members present, date and time of day were sampled during 30-min or 1-hr focal follows. The female’s behaviour (classified as feeding, moving, standing, lying, ruminating) was sampled at 1-min intervals, and, if feeding, the vegetation type she was foraging in. Vegetation types were categorised into ∼12 major types based on the map by Ferreira (1970).

The data are provided in the four online Supplementary Information files [Rum 1993 dataset; Rum 2003 dataset; Rum 2003 habitat use; Rum 2006 dataset].

### Statistical analyses

To test the main hypothesis, we compare time spent feeding by lactating and dry females across months during the spring lactation period (January to April, inclusive) and in the early summer post-lactation period when females are recovering from the costs of reproduction (May to July, inclusive). Because environmental conditions vary considerably both within and between study years, we first used general linear models with individual monthly activity time as the dependent variable and study year mean minimum temperature (°C), mean monthly daylength (hrs) and female reproductive condition (lactating versus dry) as independent variables to assess the impact of each of these factors on activity budget (research questions [12] and [2]). We then ran a four-level nested ANOVA to isolate out the effect of reproductive condition while controlling for climatic conditions individuals nested within reproductive condition, nested within month, nested within study year. Since the subsidiary analyses for dietary category, use of shelter and altitude (research questions [3] and [4]) are based on data from a single study year, we use matched pairs t-tests (matching for month) compare standard OLS linear regressions for lactating and dry females for regressed on kid age or monthly mean temperature.

*Ethical Note*. Since the studies were purely observational, there was no requirement for ethics review.

## Results

We first test the prediction that lactating females spend more time feeding than non-lactating females during the period when kids are being suckled (late January through April) (research question [1]). We then test the Altmann model prediction that feeding time should increase over time as the kid’s energy demand increases (research question [2]). Finally, we test whether lactating females compensate for the energy costs of lactation by ruminating more or foraging on richer food (research question [3]), or make more use of shelter and lower altitudes to minimise thermal costs (research question [4]).

Fig. 2 plots the mean time spent feeding and resting by lactating and dry females in each of the three study years. Table 1 gives the GLM analyses separately for the lactation (January-April) and the post-lactation (May-July) periods, with a nested ANOVA model (month within subject within reproductive status within study year) given in addition for the first of these. Fig. 3 plots the estimated marginal means for lactating and dry females during the lactation period from the nested ANOVA.

**Fig. 2.**
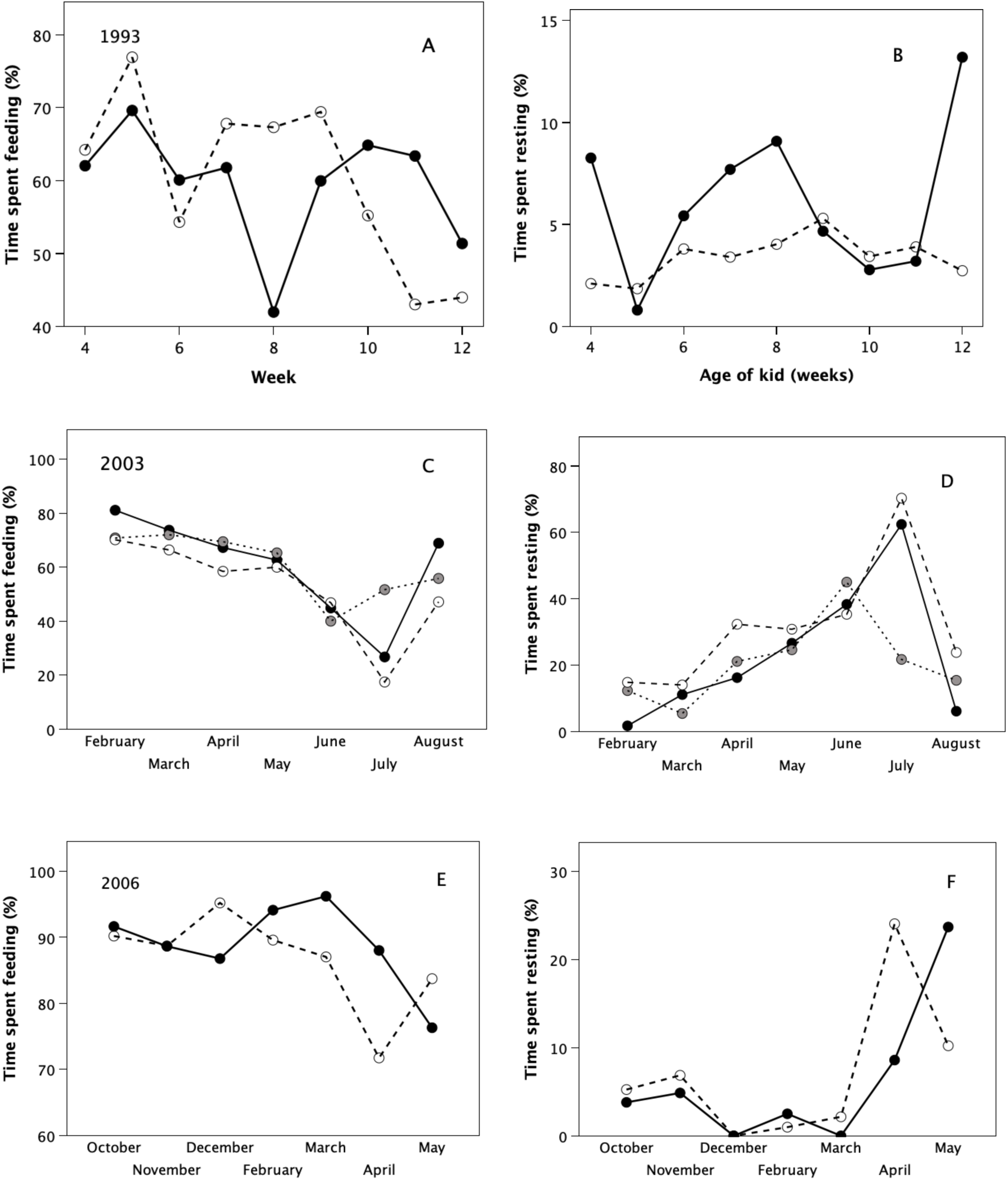
Time spent feeding and resting (lying) as a function of age of kids for the (a, b) 1993, (c,d) 2003 and (e,f) 2006 birth cohorts. Filled symbols (solid line): lactating females; grey symbols (dotted line): females who lost their kids after birth; unfilled symbols (dashed line): non-lactating (dry) females.

**Fig. 3.**
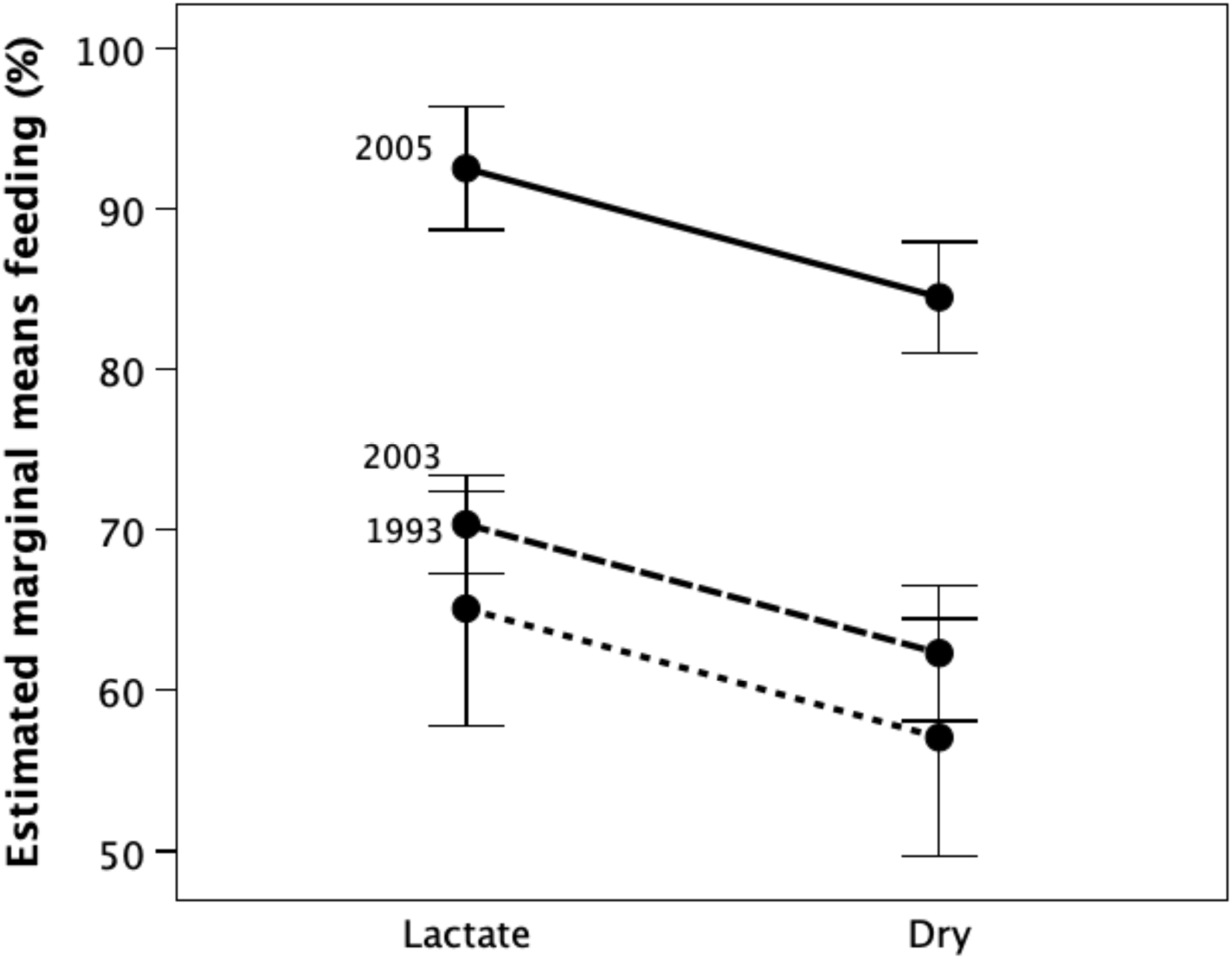
Estimated marginal means (±1 sem) for percent of time spent feeding for lactating and dry females, as a function of study year mean minimum temperature in January-April, controlling for monthly variation in daylength.

**Table 1.** Analysis of variance of feeding time (%) comparing lactating versus barren females, with study year indexed as mean minimum temperature (°C) and month indexed as mean daylength (hrs).

| Variable | F | df | p | $\eta^2$ |
| --- | --- | --- | --- | --- |
| **Lactation period (January-April)** |  |  |  |  |
| Basic full model | 5.24 | 6,53 | <0.001 |  |
| 0.372 |  |  |  |  |
| Study year: mean min temp | 14.29 | 2,53 | <0.0001 | 0.350 |
| Month: mean daylength | 4.26 | 3,53 | 0.009 | 0.194 |
| Lactating/dry | 4.50 | 1,53 | 0.039 | 0.078 |
| Nested ANOVA model* | 3.62 | 21,60 | <0.001 | 0.559 |
| Estimated marginal means: lactating = 76.0%; dry = 68.0% |  |  |  |  |
| **Post-lactation (May-June, 2003 only)** |  |  |  |  |
| Basic full model | 0.27 | 4,26 | 0.893 | 0.040 |
| Study year: mean min temp |  |  |  |  |
| Month: mean daylength | 0.19 | 3,26 | 0.817 | 0.035 |
| Lactating/dry | 0.3 | 1,26 | 0.667 | 0.007 |
| Estimated marginal means: lactating = 87.8%; dry = 89.6% |  |  |  |  |

\* month within individual subject within reproductive condition within study year

There are two key points to note. First, making due allowance for the different time scales between the three studies, the patterns are broadly similar, although feeding rates were considerably higher in 2006 than in both 1993 and 2003 (reflecting the fact that winter 2006 was significantly colder) (Fig. 3). Second, although the results were somewhat inconsistent in the 1993 study, on average during the three months of peak lactation (February through April), lactating females fed at higher rates (Fig. a,c,e), and rested at lower rates, than non-lactating females across all three studies (Fig. 2b,d,f) (Table 1). The estimated marginal means (controlling for study and month) suggest that lactating females devote ∼8 percentage points more time to feeding than dry females across the period of lactation (Table 1). In contrast, there is no significant difference between lactating and dry females during the summer months post-weaning (Table 1).

Fig. 4 plots the mean number of hours a day spent feeding (adjusted for daylength to a constant 12-hr daylength) by lactating and dry females, as well as by females whose kids had died soon after birth for the 2003 sample, along with average daylength (uppermost pecked line). The upper heavy line plots the prediction from the Altmann (1980) model (modified for goat adult and neonate body mass and kid growth rates in poor quality temperate environments from Bajhau & Kennedy 1990), benchmarked to the time that dry females spent feeding in February. Although feeding time clearly increases across months as the kids age, the slope is much shallower than that predicted by the Altmann model. After the first few weeks, lactating mothers do not come close to matching the kids’ energetic needs. Note that if lactating females were to adhere to the Altmann model, they would run out of daylight time for feeding by May. Since lactating females did not differ from dry females during the post-lactation months (and hence were not recouping their lactation energy deficit), it seems likely that they are drawing on their own body fat reserves in order to fuel lactation.

**Fig. 4.**
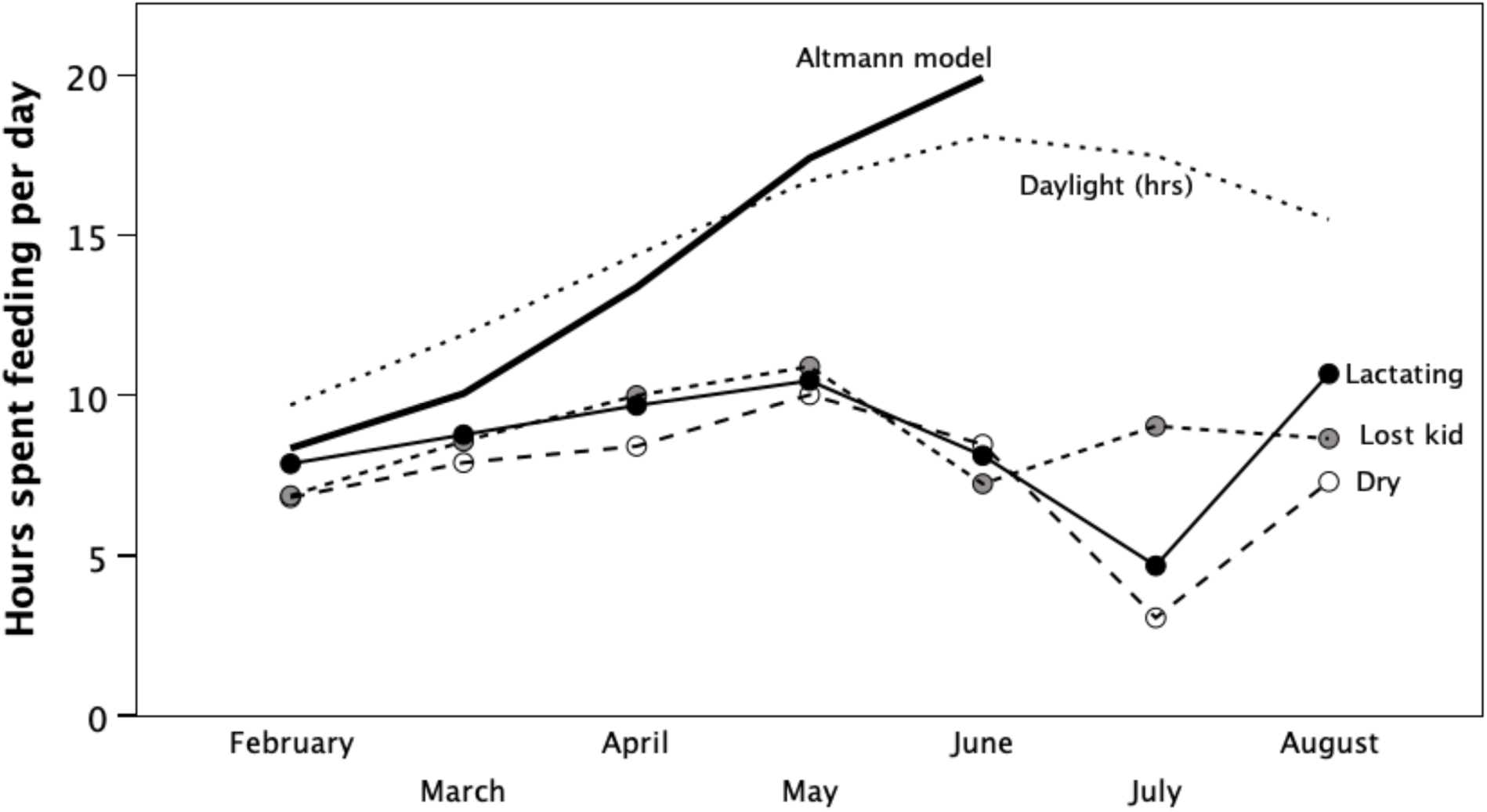
Mean number of hours per day spent feeding by non-lactating females (solid symbols, solid line), females whose kids had died shortly after birth (grey symbols, dotted line), and dry females (unfilled symbols, dashed line) over time, from the 2003 study, corrected for daylength to a constant 12-hour day. Also shown is daylength across the months (upper thin dotted line) and the feeding time for lactating females predicted by the Altmann (1980) maternal time budgets model (heavy black line). The Altmann model has been benchmarked to the observed feeding time for non-lactating females in February, with successive monthly increments based on a mean adult body mass at of 46.7kg, kid mass at birth of 2.7kg, and kid growth rate of 0.182 kg/day based on domestic goats in poor quality outdoor environments in New Zealand (Bajhau & Kennedy 1990).

That lactation might be stressful for the females is indicated by the fact that goats in New Zealand lost ∼10 kg (20% of their body mass) by the end of lactation (Bajhau & Kennedy 1990). That the Rùm females are incurring a significant cost is indicated by the fact that female mortality peaks in April shortly after kids are weaned, with a secondary peak in August after the summer birth season (Fig. 5). This pattern contrasts with male deaths, 65% of which occur between September and December, mainly as a result of the exertions incurred during the August rut (Gordon et al. 1987; Dunbar et al. 1990; Dunbar 2025). On Rùm, the bulk of the plant biomass growth occurs during the spring, such that by mid-summer the amount of nutrient-rich new growth is close to zero (Shi 2002). Females that kid in the summer face better weather conditions but significantly poorer foraging conditions than females that kid in January/February, and this probably accounts for the secondary peak in mortality in August.

**Fig. 5.**
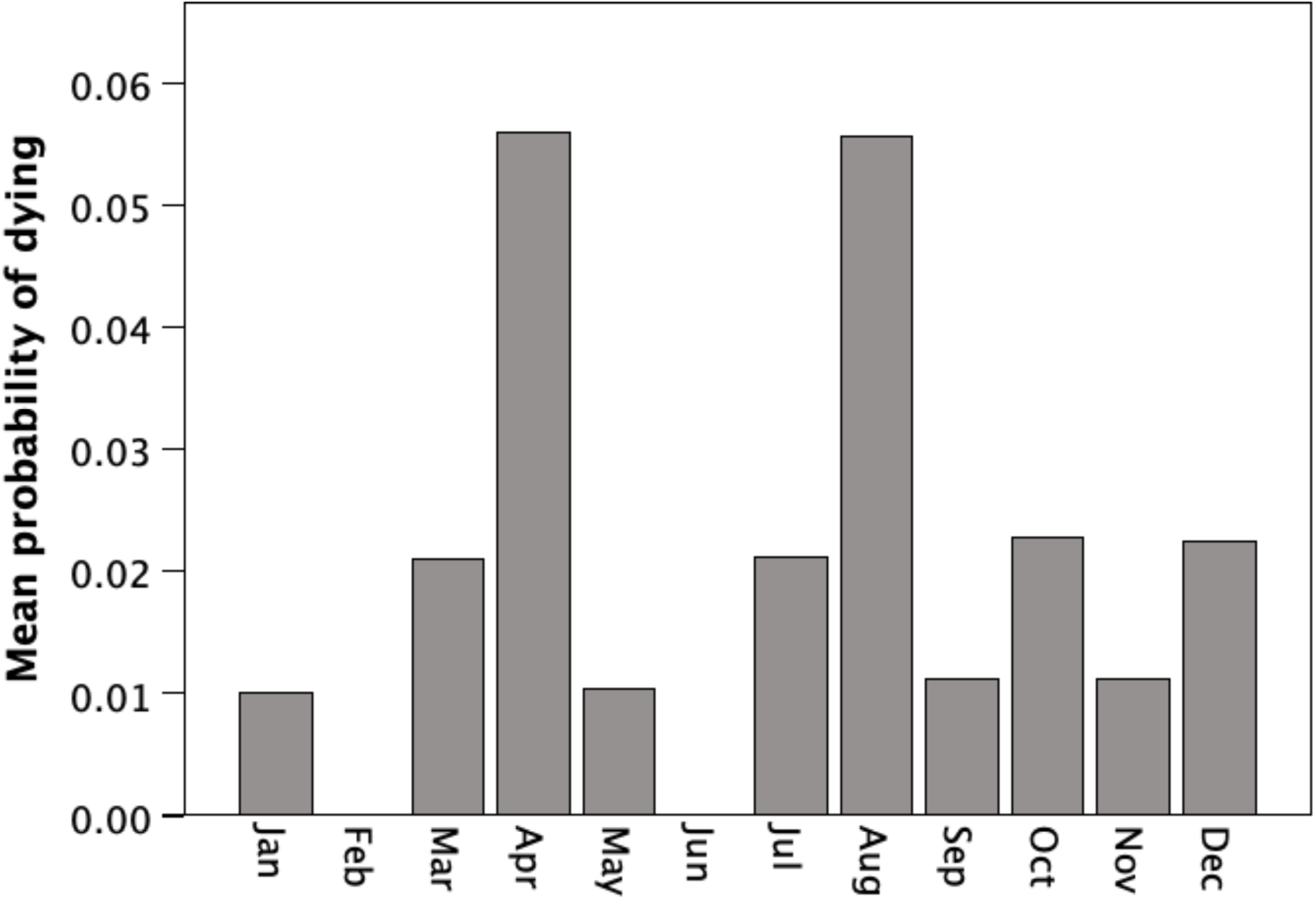
Probability of adult female mortality across the year. Data from 1981-2 study (Dunbar 2025).

To what extent can females offset the energetic costs of lactation by varying their foraging and ranging strategies so as to use their resources more efficiently and/or reduce thermal stress (research questions [3] and [4])? We considered four possibilities: (1) making use of more nutrient-rich food sources, (2) devoting more time to rumination in order to extract more nutrients from their ingesta, (3) using shelter to reduce wind chill effects, and/or (4) remaining at lower altitudes where temperatures are warmer and wind chill effects greatly reduced. Wind speeds on the cliff tops at 200m above sea level (asl) are ∼70% higher than at sea level due to wind sheer effects at ground level, which would have the effect of reducing experienced temperatures by an additional 2.5°C over ambient temperatures at the same altitude: remaining near beach rather than accompanying other goats up the cliff face during the day’s foraging would thus reduce the thermal environment by ∼4.5°C compared to the clifftops (at 200 m asl), and ∼9 °C compared to the hilltops above (at 400m asl), yielding a potentially very significant energy savings.

The relative frequencies with which females fed in the different habitat types during the 1993 and 2003 studies are shown in Fig. 6. With the exception of seaweed on the beach (t8=- 3.59, p=0.007) and *Schoenus* fen (waterlogged sedge-dominated marsh) (t8=2.54, p=0.035) in the 1993 study (lactating females avoided the first, whereas non-lactating females seemingly avoided the second), there were no significant differences in the time spent foraging in each habitat type by the lactating females compared to non-lactating females (paired-samples t-tests:1993: t8≤2.06, p≥0.073; 2003: t6≤2.15, p≥0.075). There are some differences between the two study years (e.g. greater use of herb-rich heath and grassland in 2003 than in 1993), but this may have more to do with the fact that different (albeit adjacent) heft groups were sampled than with any meaningful year-to-year differences in forage availability.

**Fig. 6.**
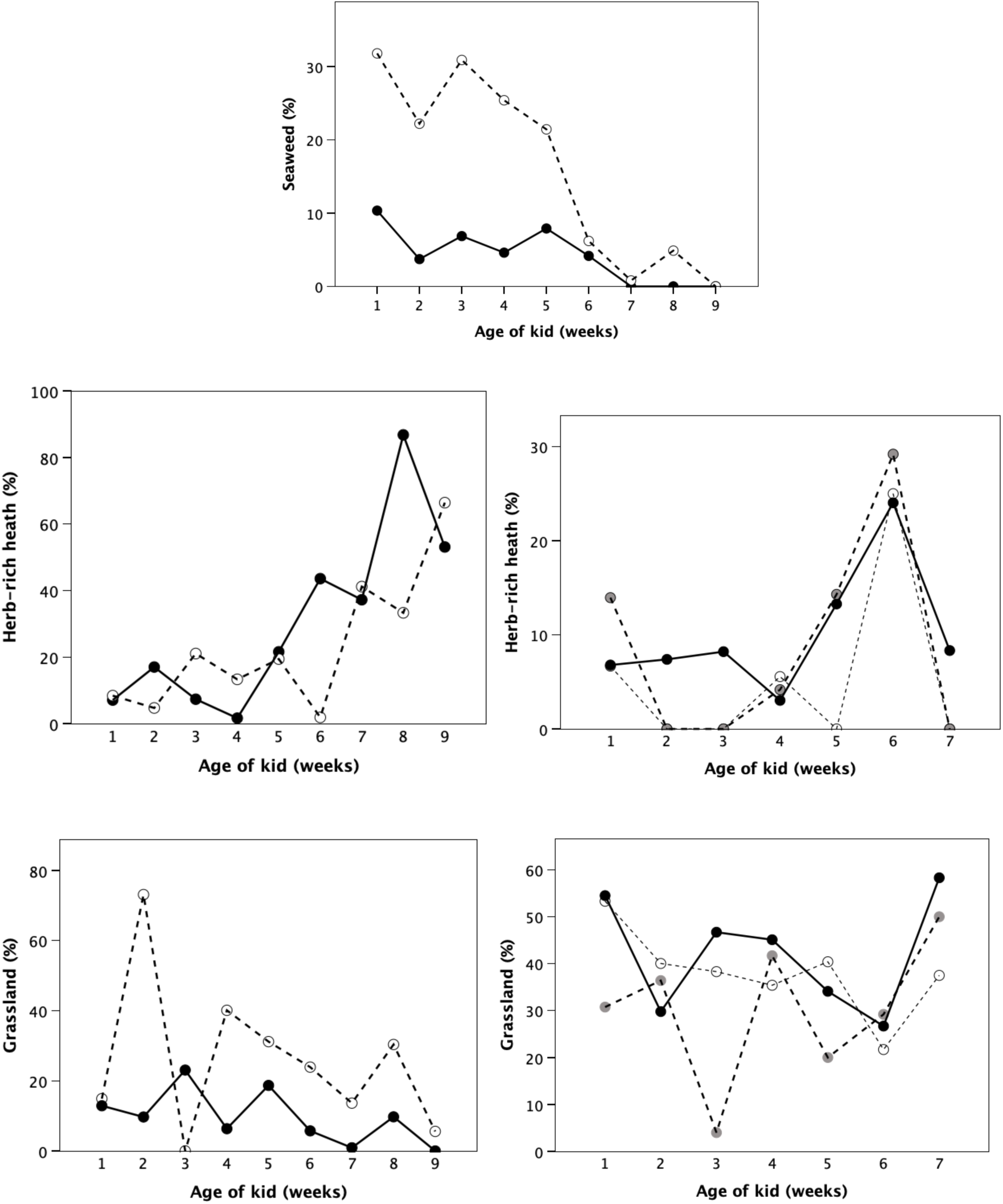
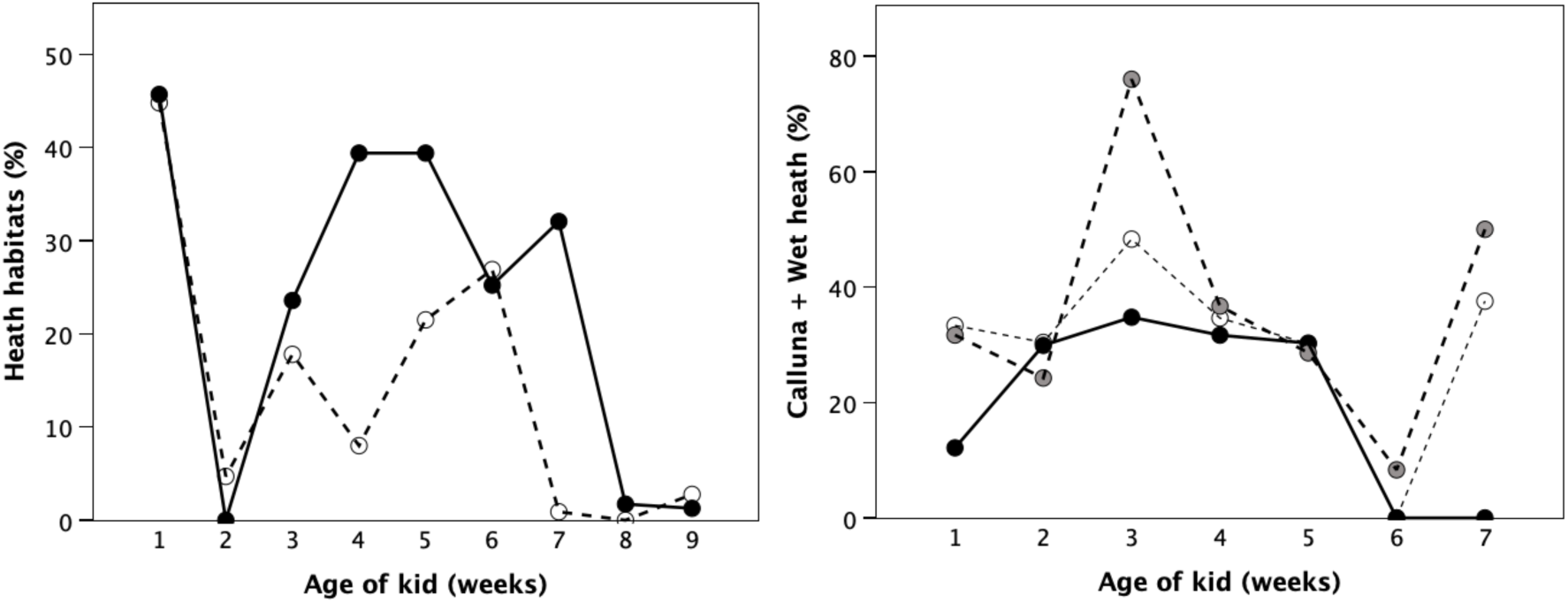
Percentage of feeding time on different food types by lactating (filled symbols, solid line) and non-lactating females (unfilled symbols, dashed lines) in successive weeks of the kid’s life in the 1993 study (a, b, d, f) and the 2003 study (c, e, g).

On balance, the dietary data suggest that lactating females were not making use of nutritionally richer food sources. However, lactating females might devote more time to rumination in order to extract more nutrients out of their ingesta. This seems not to have been the case: if anything, lactating females spent less time ruminating than other females (Fig. 7). Although, in the 2003 study, rumination rates across time for lactating females correlate with those for both females that lost their kids (rS=0.857, p=0.014) and, at least to some extent, dry females (rS=0.714, p=0.072), lactating females tended to spend less time ruminating than either of these groups of females while their kids were suckling, even though post-weaning they seemed to do more (Fig. 7).

**Fig. 7.**
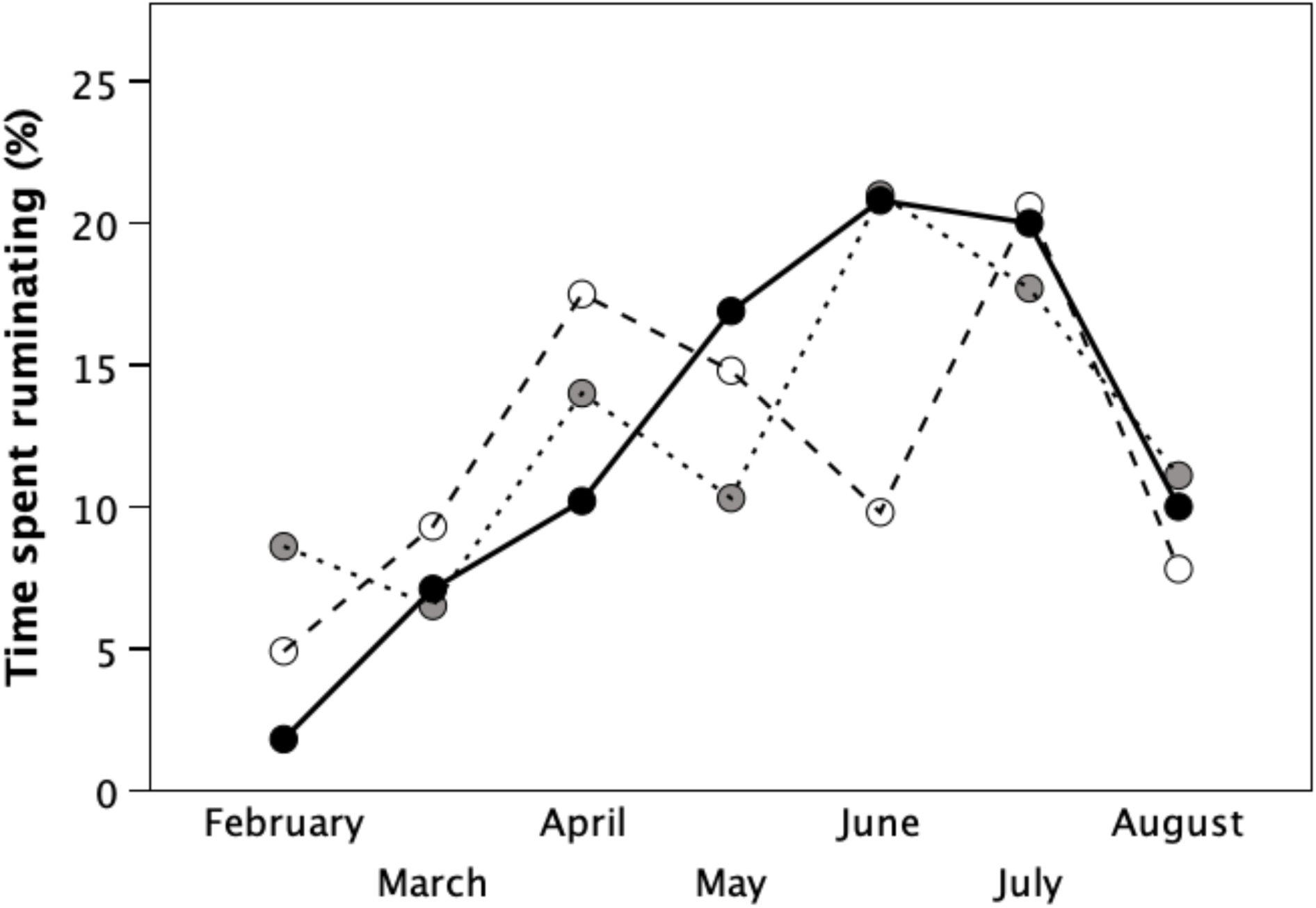
Percent of time spent ruminating (‘chewing the cud’) in each month by lactating females (filled symbols, solid line), non-lactating females (unfilled symbols, dashed line) and females whose kids had died within a few weeks of birth (grey symbols, dotted line) for the 2003 study population.

An alternative possibility is that lactating females reduce their thermal costs by making more use of shelter or by remaining at lower altitudes. For this, we used data collected in the 1993 and 2003 studies, and ran multivariate regressions for each variable separately for lactating and dry females, with time from mid-January (to control for changes in daylength), mean temperature and mean wind speed as predictor variables. Overall, lactating females did not spend significantly more time in shelter than dry females (Fig. 8; means: 20.1±20.3% vs 15.4±15.9%, respectively; t54=-0.97, p=0.367), and, if anything, they spent more time at higher altitudes (Fig. 9; means: 238.3±134.6 vs 183.1±158.9 m asl, respectively; ln-transformed altitude, t54=-2.05, p=0.045). Nonetheless, Table 2 suggests that there were significant differences between lactating and dry females in the way they responded to the thermal environment. Dry females, but not lactating females, significantly reduced their use of shelter when temperatures were warmer (p=0.047 vs p=0.334 and p=0.082 vs p=0.934 for the two cohorts, respectively, controlling for date and windspeed), suggesting that lactations were trying to avoid being excessively exposed. Note that, contrary to expectation, lactating females in the 2003 cohort spent significantly less time in shelter on windier days, despite the fact that it was windier in 2003 than in 1993 (mean Beaufort = 3.71, range 1-6, vs mean = 2.65, range 1-7, respectively). In contrast, lactating females, but not dry females, spent significantly more time at low altitudes on windier days (p=0.043 vs p=0.545, controlling for date and ambient temperature). In other words, although lactating females ranged as widely as dry females overall, they were more sensitive in the way they adjusted their exposure to the thermal environment. This would have yielded modest but significant savings in energy demand.

**Fig. 8.**
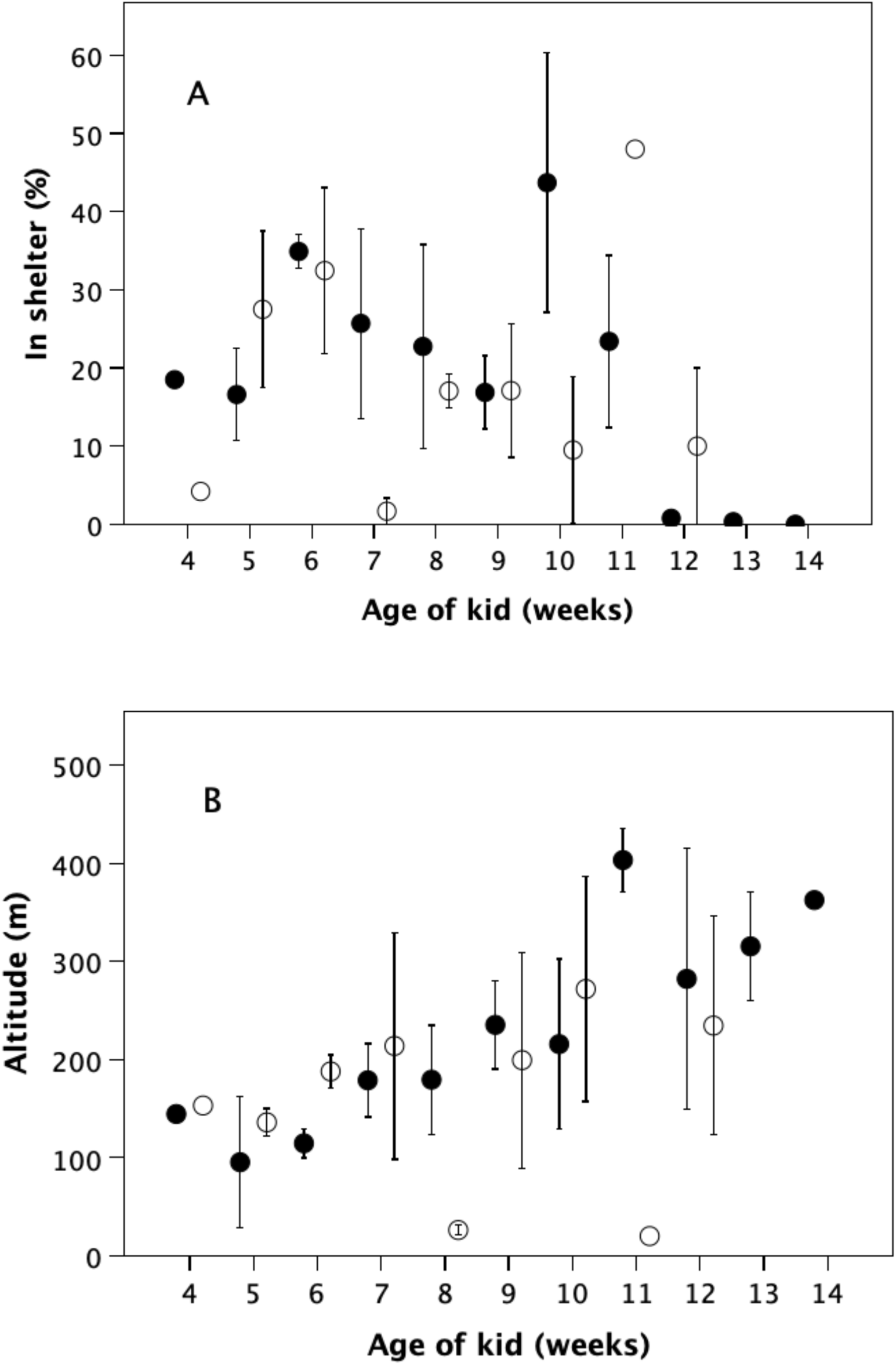
Mean (±1 s.e.m.) time spent (a) in shelter from wind and (b) at different altitudes by lactating (filled symbols) and non-lactating (unfilled symbols) females, as a function of their kid’s age, for the 1993 study population.

**Fig. 9.**
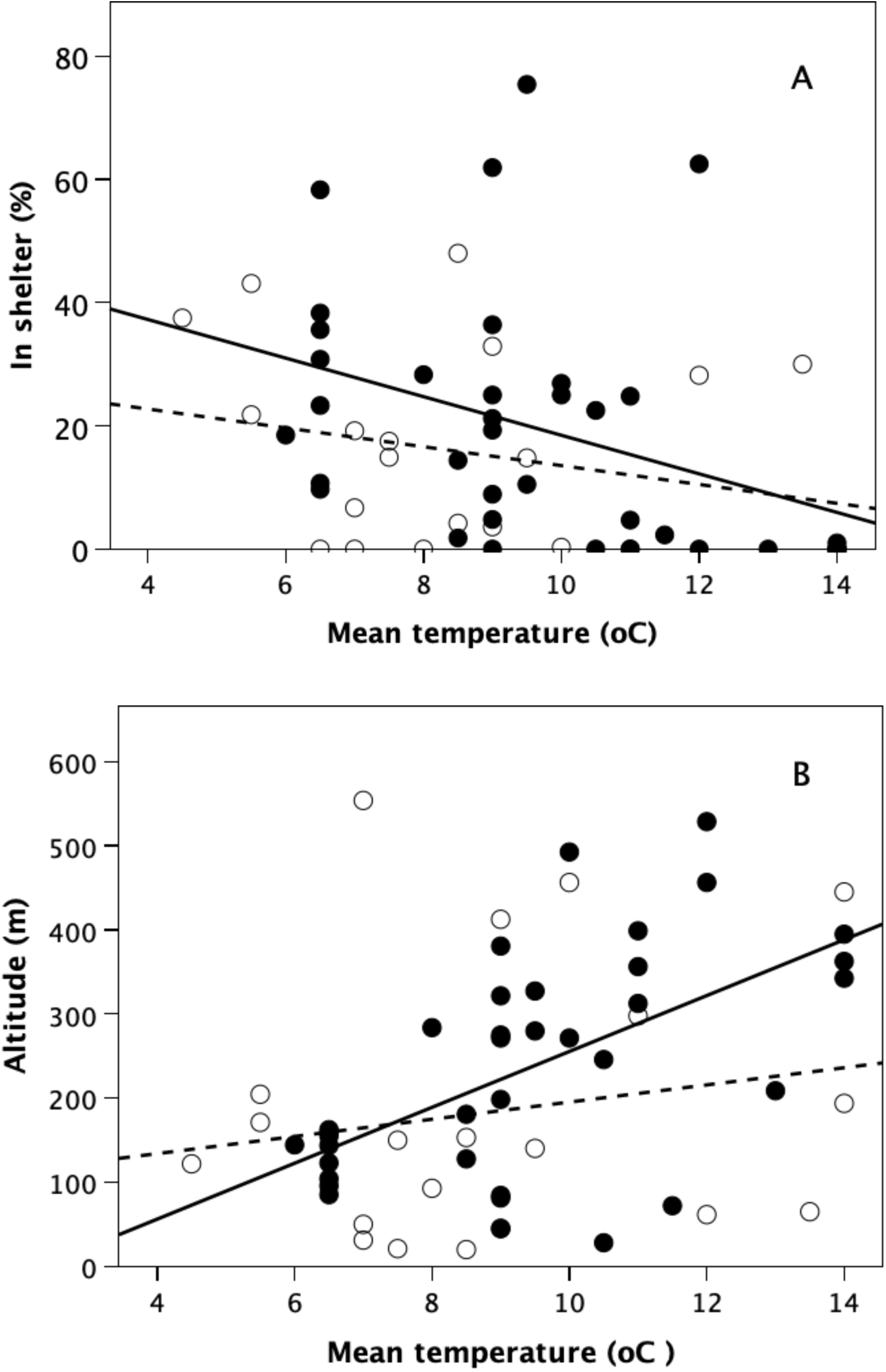
Mean time spent (a) in shelter and (b) at different altitudes by lactating (filled symbols, solid line) and non-lactating females (unfilled symbols, dashed line) in each week of the kid’s life, plotted against mean temperature that week at the study site, for the 1993 study population.

**Table 2.** Statistical analyses for the 1993 study data in Figs. 8 and 9.

| Status | predictor | slope | t | df | p |
| --- | --- | --- | --- | --- | --- |
| **Dependent = shelter (% time) (1993)** |  |  |  |  |  |
| Lactating | full model | | $F_{3,31} = 1.53$ | | 0.227 |
|  | week* | -0.432 | -0.17 | 31 | 0.867 |
|  | temperature† | -2.793 | -0.98 | 31 | 0.334 |
|  | wind‡ | -0.795 | -0.27 | 31 | 0.791 |
| Dry | full model | | $F_{3,17} = 1.58$ | | 0.232 |
|  | week* | 5.146 | 1.77 | 17 | 0.094 |
|  | temperature† | -5.107 | -2.14 | 17 | 0.047 |
|  | wind‡ | 2.681 | 1.01 | 17 | 0.329 |
| **Dependent = shelter (% time) (2003)** |  |  |  |  |  |
| Lactating | full model | | $F_{3,28} = 7.58$ | | 0.0007 |
|  | month* | -27.739 | -0.68 | 28 | 0.504 |
|  | temperature† | 2.242 | 0.08 | 28 | 0.934 |
|  | wind‡ | -19.095 | -2.24 | 28 | 0.034 |
| Dry | full model | | $F_{3,8} = 5.52$ | | 0.024 |
|  | month* | 103.054 | 1.54 | 8 | 0.162 |
|  | temperature† | -97.400 | -1.99 | 8 | 0.082 |
|  | wind‡ | 1.450 | 0.09 | 8 | 0.928 |
| **Dependent = ln(mean altitude, m asl) (1993)** |  |  |  |  |  |
| Lactating | full model | | $F_{19,36} = 6.02$ | | 0.002 |
|  | week* | 0.127 | 1.66 | 31 | 0.108 |
|  | temperature† | -0.002 | -0.03 | 31 | 0.978 |
|  | wind‡ | -0.187 | -2.11 | 31 | 0.043 |
| Dry | full model | | $F_{3,17} = 0.79$ | | 0.514 |
|  | week* | -0.267 | -1.39 | 17 | 0.182 |
|  | temperature† | -0.002 | -0.03 | 17 | 0.144 |
|  | wind‡ | -0.187. | -2.11 | 17 | 0.545 |
| Source: 1993 study |  |  |  |  |  |
| * week or month from start of birth season (mid-January); † mean weekly °C; |  |  |  |  |  |
| ‡ mean Beaufort wind speed scale [analogue scale 0-13] |  |  |  |  |  |

## Discussion

As with larger brained primates, lactating female goats spent more time foraging than non-lactating females, but the increase was considerably lower (rarely more than an extra 5-8 percentage points) and of very much shorter duration (6-8 weeks compared to 6-8 months for catarrhine monkeys and apes) than is the case for baboon mothers. Although ungulates bear the cost of lactation for only a short period of time, the extra feeding they did was unlikely to have offset the energetic costs they incurred. Lactating females did not seem to forage in ways that are significantly different from non-lactating females, making it unlikely that these females were able to exploit richer food sources to pay for the costs of lactation. They did, however, make more nuanced use of shelter and lower altitudes when thermal conditions were adverse, which will certainly have helped reduce thermoregulatory costs in this thermally challenging environment, especially for the kids (whose energy costs the females have to provide for by lactation). With ambient temperatures at beach level ∼4°C warmer than on the hilltops (Dunbar & Shi 2013), and an additional 2.5°C saving on wind chill effects, the net energetic benefit is likely to have been considerable. However, the seasonal pattern of mortality (Fig. 5) suggests that females were unable to fully compensate for the energetic costs of lactation, with the result that some failed to survive through the summer.

Ruckstuhl & Festa-Bianchet (1998) found that lactating bighorn sheep ewes did not spend significantly more time foraging during the summer period when their lambs were suckling compared to non-lactating females, but they did increase both the time spent foraging and bite rate during the autumn following weaning (see also Ruckstuhl et al. 2003), presumably to replenish fat reserves. They suggested that the ewes compensated for the body mass lost during lactation by not reducing their feeding rates during the autumn in the way that non-lactating ewes did. In contrast, lactating female alpine ibex spent significantly more time grazing than non-lactating females, and fed more quickly (higher bite rates) (Neuhaus & Ruckstuhl 2002). Lactating and non-lactating females of the Cantabrian chamois (*Rupicapra pyrenaica*), however, do not seem to differ, suggesting that they do not or cannot increase forage intake (Pérez-Barberia & Nores 1996). Other strategies are, of course, possible: female grey seals typically store up fat reserves during the non-breeding season to meet the very high energy demands of lactation (Fedak & Anderson 1982). There is some suggestion that the Rùm goats devote more time to feeding during the summer months in a way that would allow them to replenish body reserves and build up fat stores for pregnancy during the coming autumn (Dunbar & Shi 2013). Moreover, the Rùm females typically gave birth only in alternate years (as did most of the sympatric red deer hinds), suggesting that recovery from reproduction took an unusually long time compared to lower latitude populations (most of which twin every year).

It is possible that food processing constraints may influence the goats’ feeding time. In ungulates, forage intake, and therefore foraging time, is a direct function of the speed at which plant tissue can be digested (Spalinger et al 1988), with digestive capacity limited by stomach size and rumen turnover rates (Robbins 1983; van Soest 1994). Incisor bar size (Gordon & Illius 1988; Gordon et al. 1996) and plant structure (Illius & Gordon 1993) also restrict forage intake.

These effects may prevent goats from compensating fully by increasing the rate of plant matter consumption during the lactation period. Lactating deer hinds (which are much larger) adjusted their diet to feed more on grasses and less on heather compared to yeld (dry) hinds (Blaxter *et al* 1974). Although forage choice by the goats certainly changed over the course of lactation, much of this can probably be attributed to differences in availability rather than preference. However, the main issue for the lactating goats may have been that they were restricted in their ability to seek out high quality food sources by their kids’ limited mobility in a physically and thermally very challenging landscape.

Feral goat populations at high latitudes face two major constraints to which, as a mid-latitude species, they are not especially well adapted: short winter daylengths combined with a very stressful thermal environment. Not only does this impose considerable stress on non-reproductive animals, but the costs of lactation add a significant extra burden for reproductive females. Female goats on Rùm did not seem to have much leeway in which to draw on surplus time (especially during the late winter lactation period) or other adaptations to cope with these pressures. Instead, it seems that they had to draw on their own body fat reserves and then try to recoup these losses in the late spring once their kids had been weaned. Many, it seems, fail. This suggests that species invading temperate and Holarctic environments from the tropics face very considerable selection pressures and are forced to undergo relatively rapid adaptation in order to successfully occupy habitats that, at the latitude of Rùm, are close to the limits of the goats’ physiological tolerances (Dunbar & Shi 2013).

The failure of the goats to match the Altmann (1980) model has important implications in relation to offspring growth rates by comparison with primates. Goat kids are heavier at birth than primate infants, and grow and mature much faster. They therefore require heavy maternal investment. Goats are effectively adult by 12 months of age (though few females reproduce before the age of two years); large-bodied Old World monkeys and apes do not achieve puberty (never mind full physical adulthood) until they are 3-8 years old. To achieve this, primates have to spread the costs of maternal investment in lactation over a much longer period than goats do for the reasons implied by Fig. 4: they would very quickly run out of foraging time if they tried to produce an energetically expensive large-brained offspring. It seems that the only way this can be done is to slow down the rate of growth, just a anthropoid primates do. The typical large-bodied baboon weighs in at only 0.775 kg at birth and its rate of growth through to weaning averages around 0.005 kg a day (Altmann 1980); goat kids weigh 2.7 kg at birth, and grow at a rate of 0.182 kg/day (Bajhau & Kennedy 1990). In other words, goat kids grow ∼35 times faster than baboons, despite their larger size. This constraint has significant implications for ungulates’ capacity to grow large-brained offspring: without a major change of dietary strategy, it would be impossible for them to do so.

## Acknowledgments

We thank the research assistants who helped with data collection (Katie Stirling in 1993; Annie Harrison and Ian MacDonald in 2006), Scottish Natural Heritage for permission to work on Rùm, and the Wardens and staff of the Isle of Rùm Nature Reserve for their continued support over the years.

## Statement of Ethics

The research reported herein was purely observational and did not require ethical approval.

